# SARS-CoV-2 infection reduces Krüppel-Like Factor 2 in human lung autopsy

**DOI:** 10.1101/2021.01.15.426691

**Authors:** Tzu-Han Lee, David Wu, Robert Guzy, Nathan Schoettler, Ayodeji Adegunsoye, Jeffrey Mueller, Aliya Hussein, Anne Sperling, Gökhan M. Mutlu, Yun Fang

## Abstract

Acute respiratory distress syndrome (ARDS) occurred in ~12% of hospitalized COVID-19 patients in a recent New York City cohort. Pulmonary endothelial dysfunction, characterized by increased expression of inflammatory genes and increased monolayer permeability, is a major component of ARDS. Vascular leak results in parenchymal accumulation of leukocytes, protein, and extravascular water, leading to pulmonary edema, ischemia, and activation of coagulation associated with COVID-19. Endothelial inflammation further contributes to uncontrolled cytokine storm in ARDS. We have recently demonstrated that Krüppel-like factor 2 (KLF2), a transcription factor which promotes endothelial quiescence and monolayer integrity, is significantly reduced in experimental models of ARDS. Lung inflammation and high-tidal volume ventilation result in reduced KLF2, leading to pulmonary endothelial dysfunction and acute lung injury. Mechanistically, we found that KLF2 is a potent transcriptional activator of Rap guanine nucleotide exchange factor 3 (RAPGEF3) which orchestrates and maintains vascular integrity. Moreover, KLF2 regulates multiple genome-wide association study (GWAS)-implicated ARDS genes. Whether lung KLF2 is regulated by SARS-CoV-2 infection is unknown. Here we report that endothelial KLF2 is significantly reduced in human lung autopsies from COVID-19 patients, which supports that ARDS due to SARS-CoV-2 is a vascular phenotype possibly attributed to KLF2 down-regulation. We provide additional data demonstrating that KLF2 is down-regulated in SARS-CoV infection in mice.

## Introduction

ARDS occurred in ~12% of hospitalized COVID-19 patients in a recent New York City cohort and has a high mortality rate depending on age decile (1). Pulmonary endothelial dysfunction, characterized by increased expression of inflammatory genes and increased monolayer permeability, is a major component of ARDS. Vascular leak results in parenchymal accumulation of leukocytes, protein, and extravascular water, leading to pulmonary edema, ischemia, and activation of coagulation associated with COVID-19. Endothelial inflammation further contributes to uncontrolled cytokine storm in ARDS (2). Recent studies reported endothelial dysfunction in lung autopsy of COVID-19 patients (3, 4).

We have recently demonstrated that Krüppel-like factor 2 (KLF2), a transcription factor which promotes endothelial quiescence and monolayer integrity (5), is significantly reduced in experimental models of ARDS. Lung inflammation induced by influenza A H1N1 virus or lipopolysaccharide (LPS) leads to reduced KLF2, which drives pulmonary endothelial dysfunction and acute lung injury (6). High-tidal volume ventilation in rat mimicking ventilator-induced lung injury (VILI) also reduces KLF2 and drives lung injury (6). Mechanistically, we found that KLF2 is a potent transcriptional activator of Rap guanine nucleotide exchange factor 3 (RAPGEF3), which activates Ras-related C3 botulinum toxin substrate 1 (Rac1) (6), a key small GTPase that orchestrates and maintains vascular integrity by stabilizing cortical actin. Moreover, KLF2 regulates multiple genome-wide association study (GWAS)-implicated ARDS genes related to cytokine storm, oxidation, and coagulation in lung microvascular endothelium (6).

Whether lung KLF2 is regulated by SARS-CoV-2 infection is unknown. Here we report that endothelial KLF2 is significantly reduced in human lung autopsies from COVID-19 patients, which supports that ARDS due to SARS-CoV-2 is a vascular phenotype possibly attributed to KLF2 down-regulation. We provide additional data demonstrating that KLF2 is down-regulated in SARS-CoV infection in mice.

## Methods

Within 24-72 hours of death, autopsies of COVID-19 patients were performed and lung tissue was stored in 10% neutral buffered formalin for at least 72 hours per University of Chicago protocol. Lung tissue from COVID-19 donors were randomly selected and compared to control lungs which were obtained from organ donors whose lungs were non-transplantable, and without SARS-CoV-2 infection, collected prior to the pandemic. Control lungs (obtained from Gift of Hope, GOH) were similarly processed between 24-72 hours of death and stored in 10% neutral buffered formalin for at least 72 hours. Lung tissue were then embedded in paraffin and sectioned. Tissue slices were then processed with conventional hematoxylin and eosin staining or immunofluorescence. Deparaffinization and rehydration was performed by serial washes with decreasing amounts of ethanol with a final wash in water. Samples were then blocked with 1% BSA in PBS for 30 minutes followed by permeabilization in 1% BSA and 0.1% TritonX-100 for 1 hour at room temperature. Immunofluorescence was performed with overnight incubation with primary antibodies to KLF2 and vascular endothelial cadherin (VE-Cad), the latter to delineate endothelial cells. Samples were incubated with secondary antibodies anti-goat Alexa 488 and anti-mouse Alexa 647 for 1 hour at room temperature. Washes were performed with PBS. A confocal microscope was used to acquire fluorescent images.

## Results

Detailed clinical information was available for all patients with COVID-19 (**Table 1**). At diagnosis, the patients had severe acute hypoxemic respiratory failure and required either simple nasal cannula, high flow nasal cannula, or mechanical ventilation, meeting Berlin criteria for ARDS. Chest X-rays showed multifocal bilateral alveolar opacities (data not shown). Pathological findings are also detailed in **Table 1**. Autopsy results in COVID-19 patients demonstrate pathological ARDS with diffuse alveolar damage as well as a significant component of superimposed pneumonia.

**Table 1.**

| Case #* | Age | Sex | Ethnicity | Smoking status | Length of hospital stay | Ventilatory support^ | P/F ratio | Lung autopsy findings |
| --- | --- | --- | --- | --- | --- | --- | --- | --- |
| C1 | 36 | M | Black | Yes | 13 | HFNC, MV | 64 | Acute bronchopneumonia and few microthrombi |
| C2 | 84 | F | Black | Former | 4 | NC | 191 | Acute and chronic aspiration pneumonia, focal bronchopneumonia |
| C3 | 84 | M | Black | N/A | 1 | HFNC | 65 | Reactive pneumocytes and fibrin thrombi with bilateral bronchopneumonia |
| C4 | 77 | F | Black | N/A | 15 | HFNC | NA | Hyaline membrane formation and focal superimposed bronchopneumonia |
| C5 | 51 | F | Black | Yes | 12 | HFNC, MV | 64 | Diffuse alveolar damage and mucous plugging |
| C6 | 82 | F | Black | Former | 13 | HFNC, MV | 63 | Diffuse alveolar damage, hemorrhage, aspiration pneumonia |
| C7 | 84 | F | Black | No | 5 | NC | N/A | Acute and chronic aspiration pneumonia |
| C8 | 40 | M | Black | No | 25 | MV | 62 | Diffuse alveolar damage, thromboemboli, areas of infarction |
| C9 | 80 | F | Black | Former | 1 | N/A | N/A | Diffuse acute pneumonia and focal platelet thrombi |
| C10 | 78 | F | Black | Former | 5 | MV | 141 | Diffuse alveolar damage and intraalveolar hemorrhage |
| R1 | 58 | M | N/A | Yes | N/A | MV | N/A | Recent hemorrhage (trauma) |
| R2 | 59 | F | N/A | Yes | N/A | MV | N/A | Patchy non-specific interstitial fibrosis |
| R3 | 53 | M | N/A | Former | N/A | MV | N/A | Emphysema |
| R4 | 35 | M | N/A | Yes | N/A | MV | N/A | Emphysema, patchy non-specific interstitial fibrosis |
| R5 | 59 | F | N/A | Yes | N/A | MV | N/A | Patchy non-specific interstitial fibrosis |

As shown in **Figure 1**, GOH control lungs (n = 5) expressed significantly higher levels of KLF2 protein in the lung compared with lungs from COVID-19 patients (n = 10). KLF2 expression largely co-localized with VE-Cad. Only KLF2 protein but not VE-Cad was reduced in the SARS-CoV-2-infected lungs, suggesting KLF2 reduction in COVID-19 patients is not due to endothelial loss. Representative immunofluorescence images with histological counterparts are provided.

**Figure 1.**
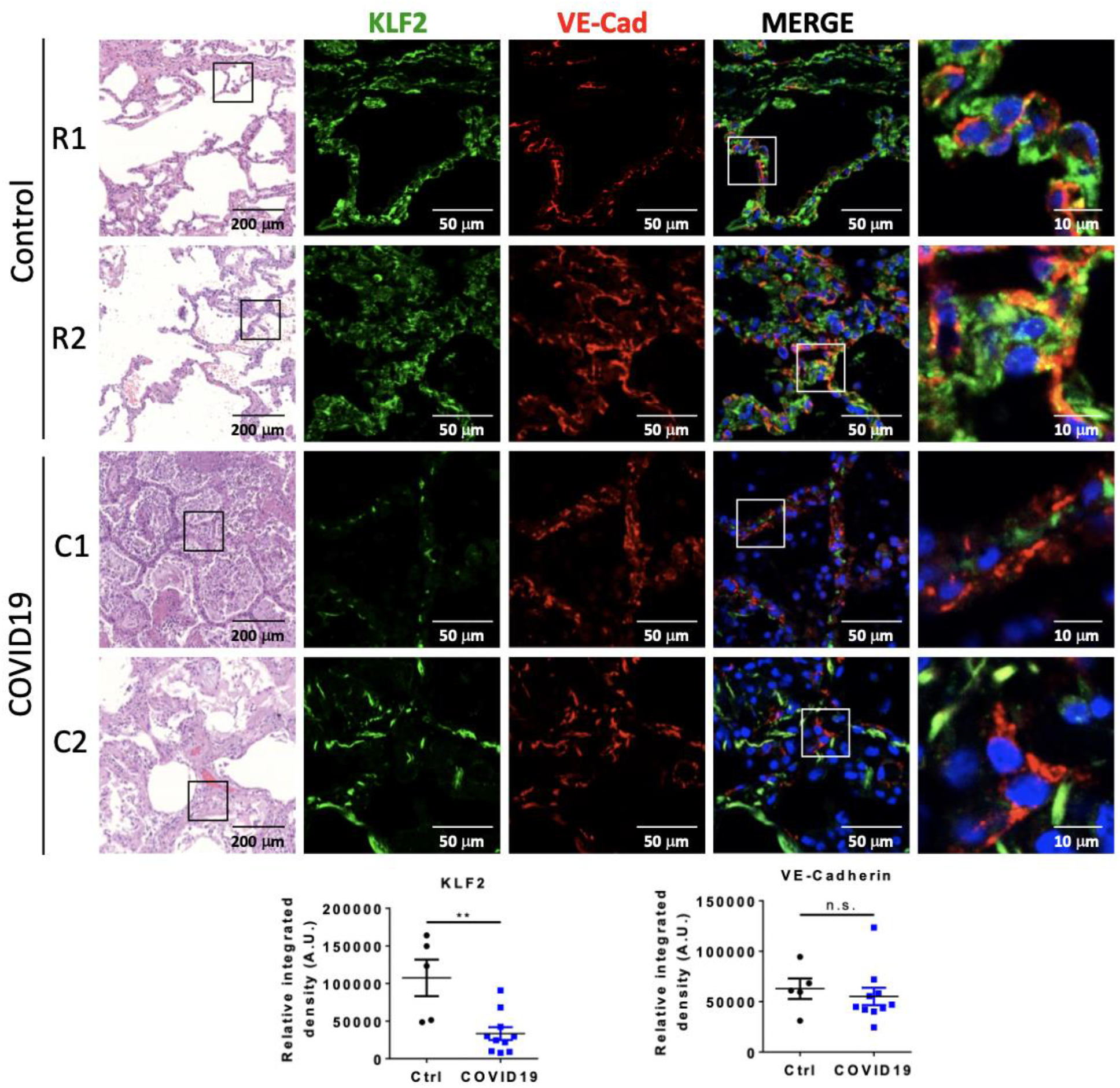
Both ROBI (R#) and COVID-19 (C#) lung autopsy samples demonstrate distorted architecture on histology. Immunofluorescence is performed for KLF2 and VE-Cad, and quantified below. In ROBI lungs, KLF2 signal (green) co-localized with VE-Cad (red), which highlights pulmonary vasculature (n = 5). In COVID-19 samples, the KLF2 signal is significantly reduced when compared to ROBI lungs (p < 0.005, n = 10). There is no significant difference in VE-Cad between ROBI and COVID19 samples.

We reanalyzed microarray results in which mice were infected intranasally with 10^5^ PFU of mouse-adapted SARS-CoV (7), showing that lung KLF2 mRNA is reduced by ~40% due to SARS-CoV infection (Mock: 16398 +/- 1722 vs SARS-CoV: 9977 +/- 844.7, p < 0.05, n=5 each). These data collectively demonstrate that endothelial KLF2 is significantly reduced in lungs infected by coronavirus, supporting recent human lung autopsy studies showing vascular inflammation and endothelial dysfunction due to SARS-CoV-2 infections (3, 4).

## Discussion

Lung microvascular dysfunction is instrumental to breakdown of the alveolar-vascular barrier, resulting in edema, neutrophils recruitment, followed by radiographic opacities, and hypoxemia due to shunt formation. Endothelial KLF2 is a major regulator of microvascular quiescence and integrity. Our data demonstrate that lung endothelial KLF2 is significantly down-regulated with SARS-CoV-2 infection, suggesting that vascular destruction contributes to COVID-19 pneumonia and ARDS.

COVID-19 has generated lay controversy because of apparent disproportionate hypoxia to work of breathing; patients may have hypoxemia but relatively little dyspnea. It is unknown whether dysfunctional pulmonary blood flow itself can cause hypoxia in COVID-19 pneumonia. One proposed mechanism is dysregulated nitric oxide synthesis, which under normal conditions provides appropriate ventilation/perfusion matching. Capillary flow maintains expression of endothelial KLF2 (6), which is a direct transcriptional activator of endothelial nitric oxide synthase (eNOS/NOS3). Thus, poor blood flow in pneumonia may itself reduce nitric oxide release, independent of alveolar filling or presence of virus. KLF2 also maintains blood fluidity by inhibiting vascular thrombosis (8), another postulated mechanism for hypoxia in COVID-19. Interestingly, statin, a potent activator of endothelial KLF2, is associated with reduced mortality of COVID-19 patients (9).

Immune suppression via corticosteroids is now used to treat severe COVID-19 pneumonia (10). Theoretically, corticosteroids suppress immune cell-mediated cytokine storm which causes widespread organ dysfunction and hypotension. In endothelium, glucocorticoids were also shown to inhibit major pro-inflammatory cytokines and chemokines (11), which are also suppressed by KLF2. Thus, corticosteroids may also dampen vascular inflammation associated with cytokine storm in ARDS. Further investigation into vascular dysfunction in ARDS, especially in COVID-19 may yield mechanistic insight and therapeutic benefit with vessel targeted therapy.

## Author contributions

THL performed experiments, analyzed and interpreted data, and edited the manuscript RG performed sample collection and processing, analyzed and interpreted data, and edited the manuscript

NS and AA analyzed and interpreted data and edited the manuscript

AS, JM and AH performed sample collection and processing and interpretation of data, and edited the manuscript

DW, GM, and YF designed experiments, analyzed and interpreted data, wrote and edited the manuscript

## Sources of Support

DW: NIH K99-HL145113

NS: NIH K08-HL153955

AA: NIH K23-HL146942

GM: NIH R01-ES015024, DoD-W81XWH-16-1-0711

YF: NIH R01-HL136765, R01-HL138223

## Footnotes

* C#: COVID-19 patients. R#: GOH patients

^ HFNC: high flow nasal cannula. MV: mechanical ventilation. NC: nasal cannula.

## Notes

### Competing Interest Statement

The authors have declared no competing interest.

